# First-step Mutations during Adaptation to Thermal Stress Shift the Expression of Thousands of Genes Back toward the Pre-stressed State

**DOI:** 10.1101/022905

**Authors:** Alejandra Rodríguez-Verdugo, Olivier Tenaillon, Brandon S. Gaut

## Abstract

The temporal change of phenotypes during the adaptive process remain largely unexplored, as do the genetic changes that affect these phenotypic changes. Here we focused on three mutations that rose to high frequency in the early stages of adaptation within 12 *Escherichia coli* populations subjected to thermal stress (42°C). All of the mutations were in the *rpoB* gene, which encodes the RNA polymerase beta subunit. For each mutation, we measured the growth curves and gene expression (mRNAseq) of clones at 42°C. We also compared growth and gene expression to their ancestor under unstressed (37°C) and stressed conditions (42°C). Each of the three mutations changed the expression of hundreds of genes and conferred large fitness advantages, apparently through the restoration of global gene expression from the stressed towards the pre-stressed state. Finally, we compared the phenotypic characteristics of one mutant, *I572L*, to two high-temperature adapted clones that have this mutation plus additional background mutations. The background mutations increased fitness, but they did not substantially change gene expression. We conclude that early mutations in a global transcriptional regulator cause extensive changes in gene expression, many of which are likely under positive selection for their effect in restoring the pre-stress physiology.

## Introduction

Organisms are often exposed to stressful environments. To cope, they have evolved responses based on the duration of the stress. For example, bacteria that are exposed to increased temperatures display a transient heat-shock response, which involves up-regulation of genes encoding heat stress-proteins (Richter et al. 2010), followed by a period of phenotypic acclimation (Gunasekera et al. 2008). If the environmental stress is sustained over a long period of time, individuals may eventually accumulate mutations that result in long-term adaptation of the population. Although acute responses to stress have been well characterized (Gunasekera et al. 2008), the mechanisms of stress acclimation and adaptation are understood less well (Riehle et al. 2001; Gunasekera et al. 2008).

One possible mechanism for adaptation to stressful conditions is genetic change that produces novel traits or new physiological functions, such as antibiotic resistance or the ability to use new metabolic pathways (Blount et al. 2012; Quandt et al. 2014). Another mechanism is genetic change that leads with restoration of cellular functions to a pre-stressed state. Rather than creating new functions, restorative mutations revert some aspect of the individuals’ altered physiological state back to an unstressed or “normal” state (Carroll and Marx 2013). This pattern of restoration has been recently observed during metabolic perturbation and high-temperature adaptation (Fong et al. 2005; Carroll and Marx 2013; Sandberg et al. 2014). Although both novelty and restoration likely drive long-term adaptation, it is not yet clear which is more frequent.

Another issue that remains largely unexplored is the temporal pace of phenotypic change during adaptation to stress. After an environment becomes stressful, the acclimated state becomes the initial phenotypic state upon which natural selection acts. Thereafter, each adaptive mutation moves the population towards a phenotypic optimum (i.e. to a phenotype that best fits the stressful environment (Fisher 1930; Orr 2005)). For historical and methodological reasons (Orr 2005), most evolutionary studies have focused on the end product of adaptation, leaving the early and intermediate steps of adaptation largely unexplored.

Fortunately, studies in experimental evolution may provide insight into the sequential magnitude of fitness changes during adaptation. To date, these studies have shown that the first beneficial mutation, or the “first-step mutation”, generally confers a large fitness advantage, perhaps because early, large-effect mutations outcompete other beneficial mutations in the population. In contrast, subsequent mutations confer more moderate fitness gains (Chou et al. 2011; Khan et al. 2011), in part due to diminishing returns epistasis (Kryazhimskiy et al. 2014).

While our knowledge about the temporal trajectory of fitness continues to grow, few studies have examined shifts in phenotypes during the adaptive process. One exception is the study of Fong et al. (2005), which used microarrays to follow the phenotype of gene expression (GE) after a shift in growth from glucose to lactate and from glucose to glycerol (Fong et al. 2005). Fong et al. (2005) observed that 39% of genes were differentially expressed during the process of acclimation to glycerol medium; however, the number of differentially expressed genes decreased at an intermediate point of adaptation and also at the endpoint of the experiment. Most of the GE changes during adaptation were restorative (Fong et al. 2005)–i.e., they restored GE to normal, pre-stress levels. Unfortunately, however, the genetic bases of these phenotypic changes were not resolved; it was unclear if the changes in GE during the intermediate point of adaptation were caused by an early adaptive mutation or were caused by combinations of mutations that accumulated during the experiment. Thus, many questions regarding phenotypic adaptation remain unresolved, such as: What are the molecular effects underlying the large fitness advantage conferred by first-step mutations? And, what is the phenotypic contribution of the first-step mutation compared to later steps in an adaptive walk?

We have decided to explore these questions based on our recent, large-scale evolution experiment (Tenaillon et al. 2012). In this experiment a strain of *E. coli* B was evolved at 42°C in 114 replicate populations. After 2000 generations, we isolated single clones from each evolved population and identified the genetic changes that had accumulated during the experiment. Overall, the *rpoB* gene, which encodes the β subunit of RNA polymerase (RNAP), was the most mutated gene with a total of 87 mutations in the 114 genomes. Three *rpoB* mutations were especially interesting because they resulted in amino acid substitutions in the active site of RNAP and conferred rifampicin resistance (Rodríguez-Verdugo et al. 2013). These mutations, all of which are located in codon 572, were driven to high frequency in 12 populations and were also beneficial at high temperature in the low glucose medium (Rodríguez-Verdugo et al. 2013). Furthermore, these mutations typically appeared early in the evolution experiment. In one population, for example, the *rpoB I572L* mutation swept to fixation before 100 generations; it was furthermore shown to confer a large fitness benefit of about 20% (Rodríguez-Verdugo et al. 2013). Since these *rpoB* mutations are located in the contact region between the downstream DNA duplex and RNAP (proximal active-site), they could play key roles in modulating the enzyme’s activity in all three stages of transcription: initiation, elongation and termination (Ederth et al. 2006).

Yet, some questions remain. First, we have not identified the mechanistic bases of their fitness advantage. Second, although we know that these parallel mutations converge on having a fitness advantage at 42°C (Rodríguez-Verdugo et al. 2013), with disadvantages at low temperatures (Rodríguez-Verdugo et al. 2014), we have yet to explore if they converged on other phenotypes. That is, it is still unclear if these mutations confer the same phenotypic effects as *rpoB* mutations found in other regions of RNAP that were also selected in some lineages during the thermal adaptation experiment. In the present study, we address all these questions. Solving these questions is relevant for understanding the adaptive mechanisms prevalent in our experimental conditions and also in other evolutionary experiments, because mutations that affect transcriptional regulators (such as RNAP and the Rho termination factor) typically appear in the early stages of stress adaptation (Applebee et al. 2008; Goodarzi et al. 2009; Kishimoto et al. 2010). Numerous observations point to the possibility that highly pleiotropic mutations in transcriptional regulators could be the first step of a general mechanism of adaptation (Fong et al. 2005).

With these considerations in mind, we have investigated the timeline of adaptive change during our evolution experiment (fig. 1). To do so, we first describe growth characteristics and GE during acclimation to thermal stress at 42°C. We then explore first-step mutations in terms of their effect on growth and GE relative to the ancestral strain. More specifically, we explore whether three amino acid substitutions in the RNAP active site converge on the same expression phenotype and also whether their effects are similar to another *rpoB* mutation that affects an amino acid substitution far from the RNAP active site. Finally, we evaluated the phenotypic contribution of one of these first-step mutations relative to the end product of adaptation, as represented by two of clones from the end of the 2000 generation experiment.

**Fig. 1.**
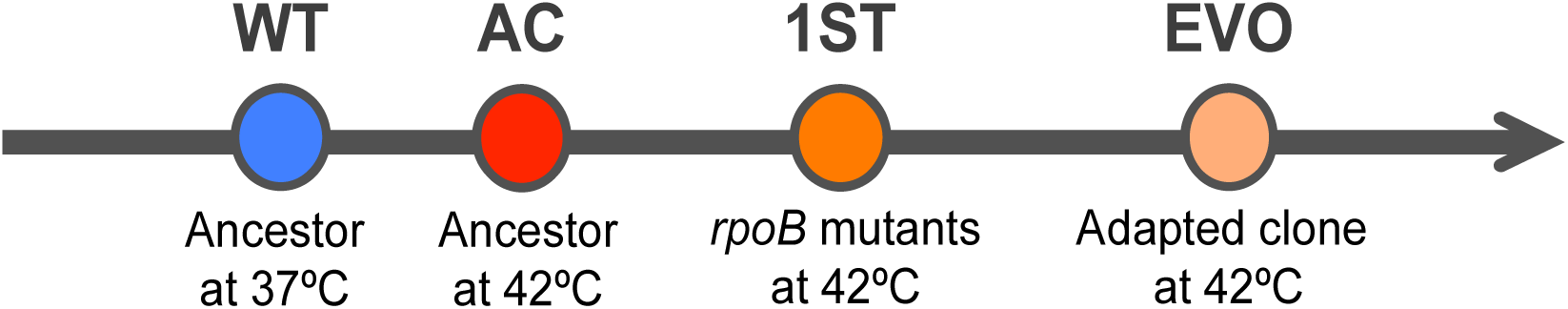
**Thermal stress adaptation timeline.** We characterized four states during thermal stress adaptation: a wild-type (WT) state, which corresponds to the ancestor grown at 37°C; an acclimatized (AC) state, which corresponds to the ancestor acclimatized at 42°C; a first-step adaptive (1^ST^) state, which corresponds to the *rpoB* single mutants; and an evolved (EVO) state, which corresponds to the high-temperature adapted clones.

## Results

### Acclimation to thermal stress involved many changes in GE

To explore the phenotypic effect of high temperature on the ancestor, we first characterized the ancestor’s growth at 37°C, which we consider to be the pre-stress condition, and at 42°C, the stress condition. The ancestor had a significantly longer lag phase and a significantly lower maximum growth rate and final yield when grown at 42°C than when grown at 37°C (fig. 2*A* and table 1), reflecting the fact that 42°C is a stressful temperature.

**Fig. 2.**
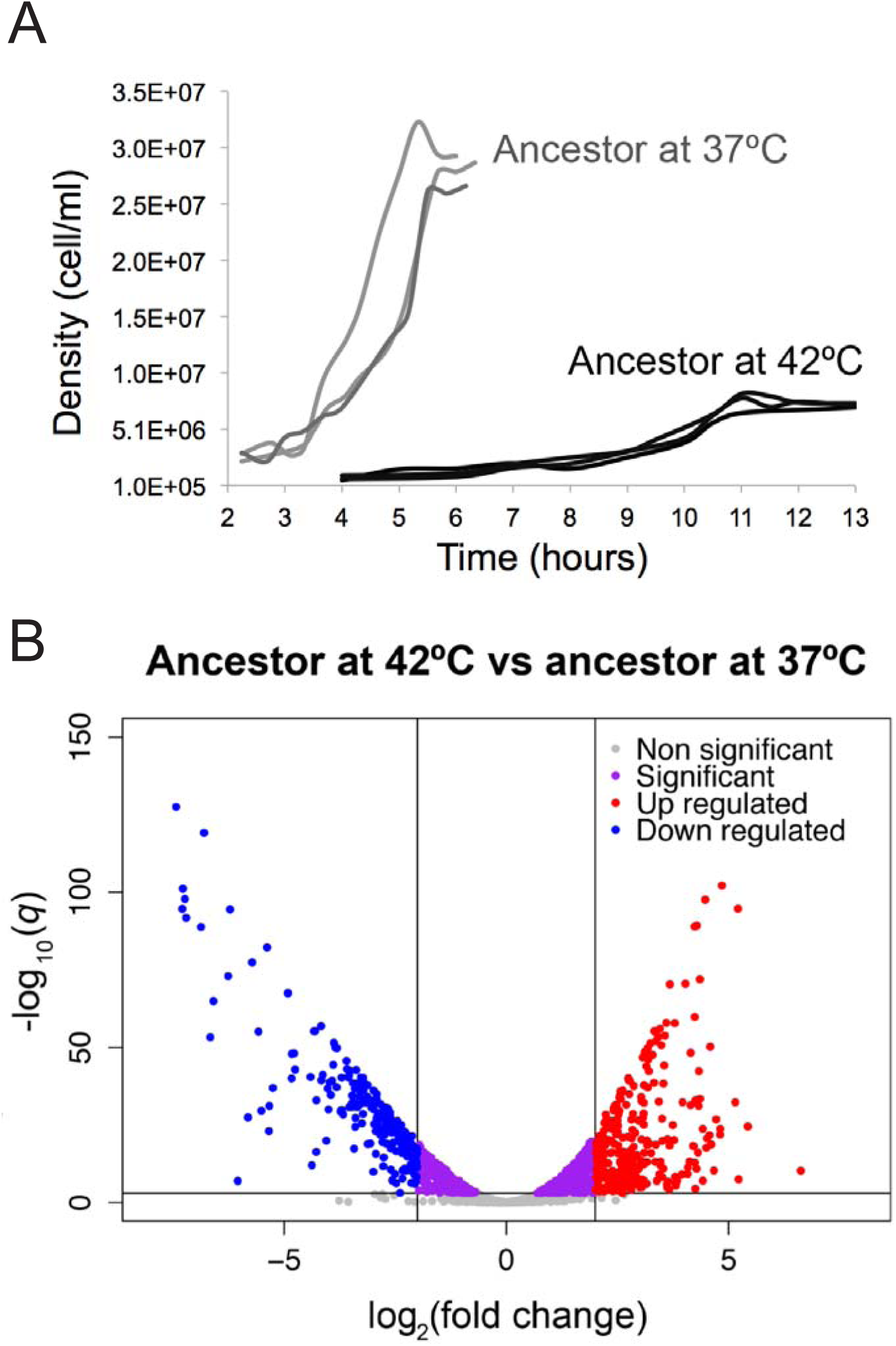
**Phenotypic characterization of the ancestor at 37°C and 42°C.** (A) Growth curves of the ancestor grown at 37°C and 42°C (three replicates at each temperature). (B) Volcano plot showing the global differential expression of genes (represented as dots) between the ancestor grown at 42°C compare to the ancestor grown at 37°C. Colors represent status with respect to 2-fold expression difference, represented by two vertical lines, and a significance at *q* = 0.001, represented by an horizontal line.

**Table 1.**
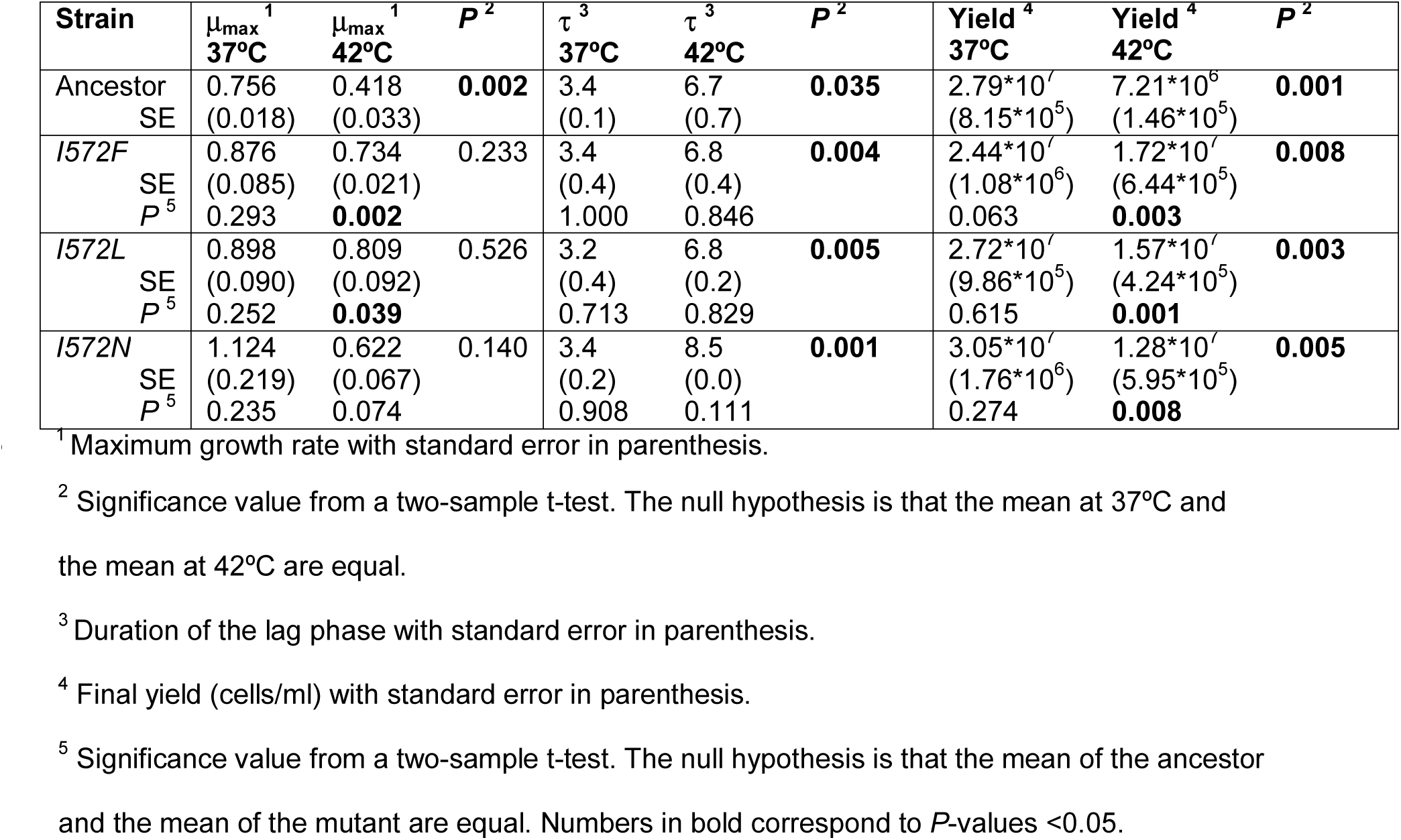
**Growth parameters of the first-step mutants and the ancestor.**

Next, we explored the global GE profile after acclimation to 42°C by obtaining RNA-seq data from three replicates of the ancestor at the mid-exponential growth phase at both 37°C and 42°C. Even under a stringent criterion of significance (*q* < 0.001), we identified 1737 genes that were differentially expressed at 42°C relative to 37°C (fig. 2*B* and supplementary dataset S1). Of these differentially expressed genes, 240 genes were highly down-regulated (log_2_fold change < -2; blue dots in fig. 2*B*) and 293 genes were highly up-regulated (log_2_fold change > 2; red dots in fig. 2*B*) at 42°C relative to 37°C. Based on an enrichment analysis of GO functional categories, we identified significant down-regulation of genes involved in translation during heat acclimation (GO: 0006412; *P* = 4.55*10^-29^). Of the genes involved in translation, 54 genes transcribed products that were structural constituents of ribosomes (GO:0003735), including *rpl*, *rpm* and *rps* genes (supplementary table S1). Other significantly down-regulated biological processes involved: *i*) amino acid biosynthesis (GO: 0008652; *P* = 2.17*10^-15^), particularly genes involved in the biosynthesis of methionine (*met* genes); *ii*) biosynthesis of ribonucleosides (GO: 0042455; *P* = 3.12*10^-7^), including purines and pyrimidine (*pur* and *pyr* genes); and *iii*) organelle organization (GO: 0006996; *P* = 2.86*10^-2^), including *flg* and *flh* genes (supplementary table S1). Surprisingly, the heat-shock inducible genes (Nonaka et al. 2006), were mostly down regulated during heat acclimation, as were the subunits of the core RNAP (*rpoA*, *rpoB*, *rpoC* and *rpoZ*) (supplementary table S2).

Among the up-regulated genes, we identified a significant enrichment of genes involved in: *i*) amino acid degradation (GO: 0009063; *P* = 6.55*10^-4^) such as degradation of arginine (*ast* genes); *ii*) alcohol degradation (GO:0046164; *P* = 5.44*10^-4^); *iii*) glycerol metabolic process (GO:0015794; *P* = 1.73*10^-2^); *iv*) monosaccharide transport (GO: 0015749; *P* = 1.11*10^-2^); and *v*) cellular response to stress (GO:0033554; *P* = 4.60*10^-2^). Of the 66 genes that had been identified previously as a component of the general stress response (Weber et al. 2005), 57 (86%) were significantly up-regulated at 42°C in our experiment (supplementary table S2).

Overall patterns of GE suggest that the ancestor at 42°C induced an early stationary phase. To be sure that this GE profile did not result from sampling at the wrong phase of the growth curve, we repeated the experiment. We again sampled three replicates at mid-exponential, repeated RNA sampling, and transcriptome analyses. The results were qualitatively similar between experiments, implying both that the ancestor at 42°C cannot unleash full growth and that the population expands at a low rate in this stressed physiological state.

### First-step adaptive mutations changed GE at 42°C dramatically

Given insights into the acclimation response at 42°C, we then investigated the effect of three potential first-step mutations – *rpoB I572F*, *I572L* and *I572N* – on cell growth and GE. Previous estimates of relative fitness indicated that these mutations are advantageous at 42°C (Rodríguez-Verdugo et al. 2013). We complemented this observation by characterizing their growth curves at both 42°C and at 37°C. The mutants had a significantly longer lag phase and lower final yield at 42°C compared to their growth at 37°C (Table 1). Nevertheless, the mutants had a significantly higher maximum growth rate and a significant higher final yield compared to the ancestor grown at 42°C, reflecting improved fitness under thermal stress (table 1 and fig. 3).

**Fig. 3.**
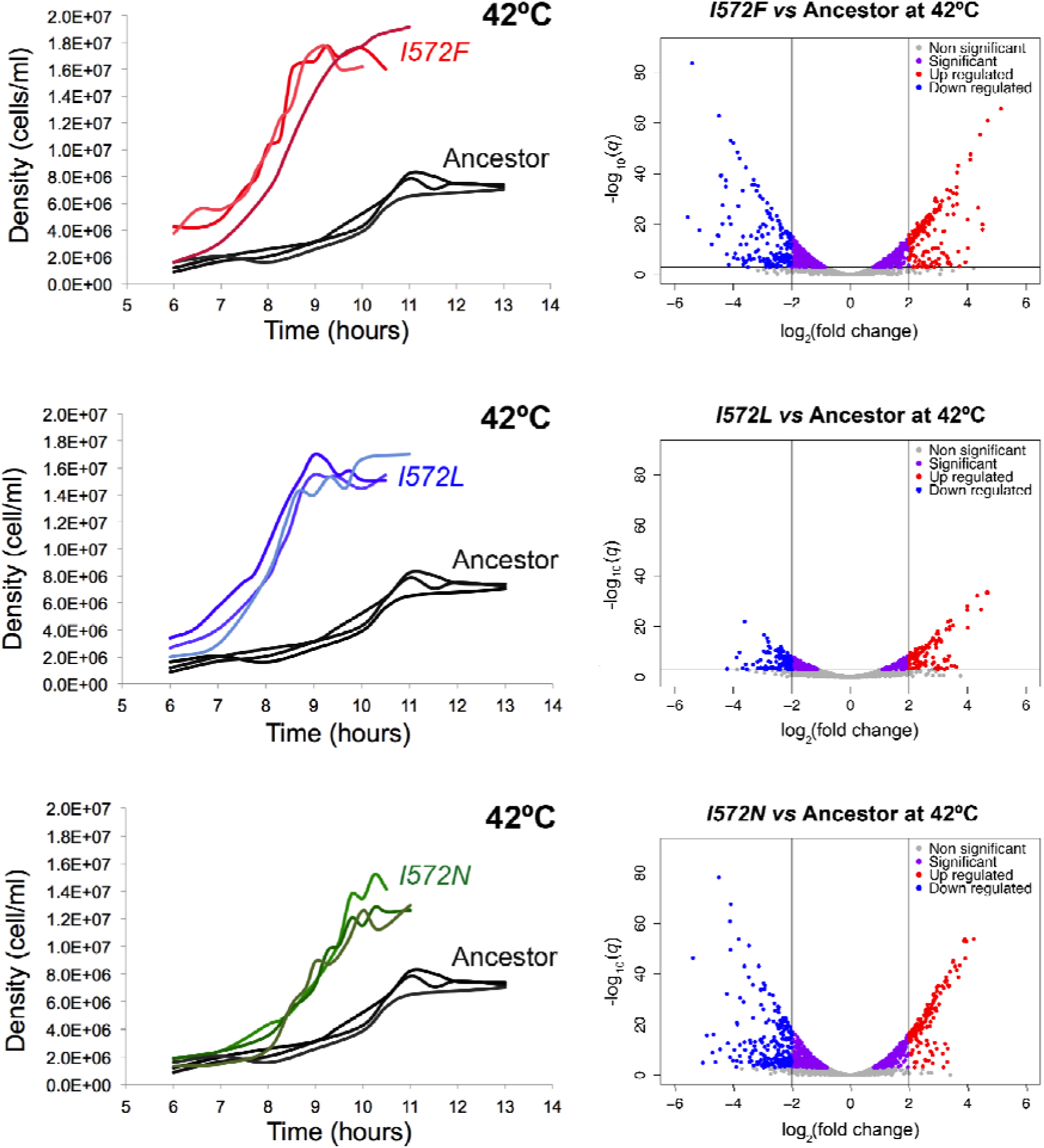
**Phenotypic characterization of the mutants compared to the ancestor at 42°C.** The left side of the figure shows the growth curves of the ancestor and the mutants grown at 42°C (three replicates per each genotype). The right side of the figure shows the volcano plots showing the global differential expression of genes (represented as dots) between the mutants grown at 42°C and the ancestor grown at 42°C. Colors represent status with respect to 2-fold expression difference, represented by two vertical lines, and a significance at *q* = 0.001, represented by an horizontal line.

We explored two hypotheses about the molecular mechanisms that may underlie the growth improvement of the mutants at 42°C. Our first hypothesis was the mutants grew better at 42°C, because the mutated RNAP was more efficient than the ancestral RNAP at transcribing DNA to RNA (Jin et al. 1992; Reynolds 2000). Our second, non-exclusive hypothesis was both that the mutated RNAP led directly or indirectly to altered transcription of a set of genes and that this differential expression underlies growth improvement (Conrad et al. 2010).

Regarding the first hypothesis, we predicted that the *rpoB* mutations slowed the RNAP complex (Rodríguez-Verdugo et al. 2014), which is otherwise accelerated at high temperatures (Ryals et al. 1982; Mejia et al. 2008). In turn, we reasoned that the reduced speed of RNAP increases both transcription fidelity and termination efficiency (Jin et al. 1988; Zhou et al. 2013), resulting in an overall higher transcription efficiency (Jin et al. 1992). To measure the transcription efficiency of the mutants and the ancestor at 42°C, we measured the relative abundance of an inducible fluorescent gene (*YFP*) inserted in our strains, at different points post induction (see *Materials and Methods*). Using this method (Reynolds 2000; Brandis et al. 2012), we found that two of the three mutants (*I572L* and *I572F*) yielded higher slopes than the ancestor at 42°C, consistent with our hypothesis of higher efficiency. However, the assay has wide confidence intervals, and there were therefore no statistical differences in the transcription efficiency of the mutants relative to the ancestor at 42°C (supplementary fig. S1). Within the limits of our experiment, we cannot conclude that the growth improvement of the mutants at 42°C is caused by higher RNAP efficiency.

We also addressed our second hypothesis that the *rpoB* mutations lead to changes in GE with consequent growth improvements at 42°C. To test this hypothesis, we obtained RNAseq data from each mutant, based on two replicates per mutant at mid-exponential growth at 42°C. We then contrasted the GE profile from each mutant against that of the ancestor grown at 42°C.

All three mutations displayed hundreds to thousand of differentially expressed genes (*q* <0.001; fig. 3). The mutant *I572F* had 1332 differentially expressed genes at 42°C with 195 highly down-regulated (log_2_fold change < -2) and 182 highly up-regulated (log_2_fold change > 2) relative to the 42°C ancestor. The mutant *I572L* had 598 differentially expressed genes with 126 highly down-regulated and 126 highly up-regulated. Finally, the mutant *I572N* had 1360 differentially expressed genes with 233 highly down-regulated genes and 164 highly up-regulated genes. To sum: single mutations within codon 572 of *rpo*B lead to numerous, pleiotropic shifts in GE.

### ***rpoB* mutations in codon 572 restored GE back toward the ancestral state.**

To investigate the general trend in GE changes from the ancestor to acclimation and then from acclimation to first-step mutations (fig. 1), we used two approaches. The first was Principal Component Analysis (PCA), which simultaneously considered the expression of all genes and all clones. The resultant plot of the first and second components exhibited differences among clones and also clearly indicated that the clones with first-step mutations shifted their gene expression profiles toward the 37°C ancestor (supplementary fig. S2).

Our second approach was to plot the log_2_fold expression change during the acclimation response (ancestor grown at 42°C *vs* ancestor grown at 37°C, x-axis) against the log_2_fold GE change during the adaptive response (mutant grown at 42°C *vs* ancestor grown at 42°C, y-axis; fig. 4). If there were no relationship between shifts in GE during acclimation and during the adaptive response, we expected a regression slope of zero; alternatively, if all the genes of a mutant were restored by an adaptive mutation – such that the expression of each genes changed from a stressed state back to a pre-stressed state – the slope of the fitted regression would approach -1.0. For the three mutants we observed a highly significant negative correlation, with slopes of -0.738 for *I572F*, -0.639 for *I572L* and -0.731 for *I572N* (fig. 4*A-C*). All three slopes were highly significant (p < 0.001) based on permutation tests (see *Material and Methods*), indicating that the main phenotypic effect of the first-step mutations was to restore global GE toward the pre-stressed state.

**Fig. 4.**
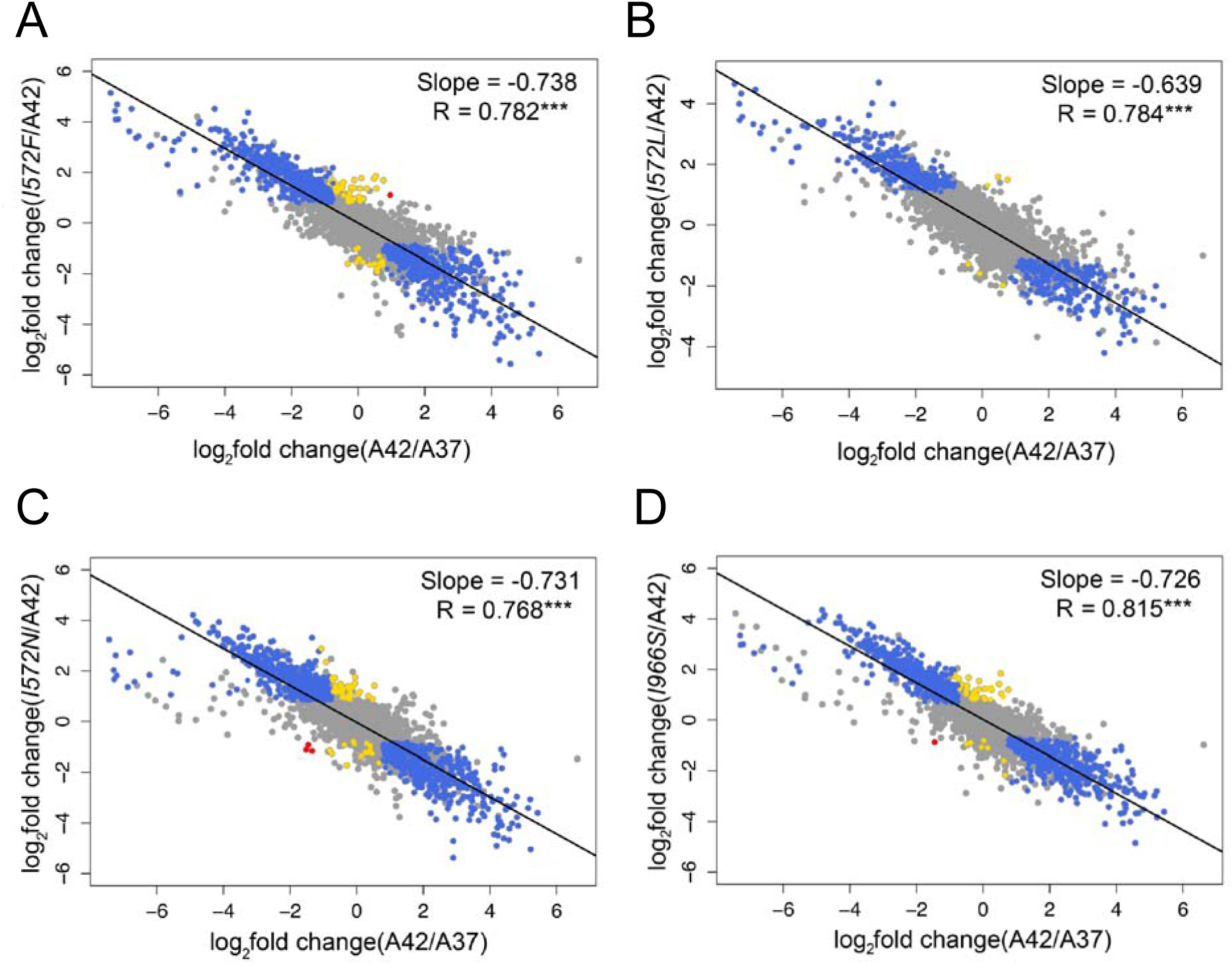
**Global changes in GE during the acclimation and adaptive response.** In all the graphs the *x*-axis represents the acclimation changes (ancestor grown at 42°C *vs* ancestor grown at 37°C). The *y*-axis represents the changes at different steps of the adaptive walk: first-step adaptive mutations (*I572F*, *I572L*, *I572N*) and a single mutant (*I966S*) *vs* ancestor grown at 42°C. Changes in expression were categorized and colored as follows: restored (blue), reinforced (red) and novel (yellow). Both unrestored and uninformative genes are colored in grey. The black line represents the linear regression fitted to the dots in each graph.

We next examined the GE changes in more detail by characterizing the expression of individual genes into one of four patterns of change that denote the direction of the effect of the mutated RNAP (see *Materials and Methods*, supplementary table S3). First, the expression of an individual gene could be *restored* back toward the pre-stressed state. Second, the expression of a gene could be *reinforced* into an exaggeration of the acclimated state, such that the mutated RNAP produced more transcripts (in the case of up-regulated genes) or fewer transcripts (in the case of down-regulated genes) than the acclimated state. Third, a gene was *unrestored* if the mutated RNAP did not change GE significantly relative to the ancestral RNAP at 42°C. Finally, GE was *novel* if it was not differentially expressed during the acclimation response, but was expressed significantly differently during the adaptive response. For these definitions, we investigated genes that differed in expression either during acclimation or between acclimation and first-mutation clones (supplementary table S3); as a result, we categorized directional shifts of roughly half of 4202 *E. coli* genes.

Following this categorization, we observed that most (63%) of the genes that were differentially expressed during acclimation were restored by mutant *I572F* (blue dots in fig. 4*A*; table 2), as expected from the strong overall negative correlation in figure 4*A*. The same was true for mutant *I572N*, for which most (67%) genes were restored (table 2). For mutant *I572L*, we found most genes to be unrestored (68%), but a substantial proportion of genes (32%) had restored expression (table 2). In contrast to restoration, only a small proportion (∼3% or less) of characterized genes exhibited novel expression in the *rpo*B mutants (table 2). While the statistical power to detect differences likely varies among clones, together the data reinforce the observation that most genes are restored in gene expression by single *rpoB* mutants.

**Table 2.** **Classification of the genes into four patterns of expression change.**

| Category | *I572F* | *I572L* | *I572N* | *I966S* |
| --- | --- | --- | --- | --- |
| 1) Restored | 1092 | 563 | 1165 | 1308 |
| 2) Reinforced | 1 | 0 | 3 | 1 |
| 3) Unrestored | 644 | 1174 | 569 | 428 |
| 4) Novel | 59 | 6 | 53 | 39 |

### **An *rpoB* mutation away from the active site of the RNAP also restores GE.**

Given that *rpoB* mutations in codon 572 converged in phenotype toward restorative GE, we sought to know if additional mutations in *rpoB* had similar effects. To address this question, we constructed a single mutant, *rpoB I966S*, which alters one of the two parallel α-helices of the Eco flap domain of RNAP (Opalka et al. 2010). The *I966S* mutation was found in 15 of our high-temperature adapted clones (Tenaillon et al. 2012), and its fitness advantage at 42°C was confirmed by competition experiments (Rodríguez-Verdugo et al. 2014).

We obtained RNAseq data from two replicates of this mutant at 42°C and performed the same GE analyses. We observed a global pattern of GE change similar to that for the *I572F*, *I572L* and *I572N* mutants. That is, the *I966S* mutation tended to restore GE from the acclimated state toward the pre-stressed state (fig. 4*D*).

To explore the convergent effects of the *I966S*, *I572F, I572L* and *I572N* mutations further, we determined the number of genes with restored GE shared among the four mutants. A third of the restored genes were shared between the four mutants, and 86% of the genes were share by at least two mutants (fig. 5), indicating a high level of expression convergence. This high level of phenotypic convergence was also highlighted by the pairwise comparisons of differential expression between the mutants (supplementary fig. S3). We performed an enrichment analysis of GO assignments for the 490 restored genes that were shared among the four mutants. Not surprisingly, given the overall pattern of restoration (fig. 4), the restored genes represented the same sets of genes that were enriched for the acclimation response. For example, significantly down-regulated genes during the acclimation response, such as the genes involved in translation (*rpl*, *rpm* and *rps* genes), were significantly up-regulated in the four mutants. This suggests that the different RNAP mutants restored GE towards a growth pattern.

**Fig. 5.**
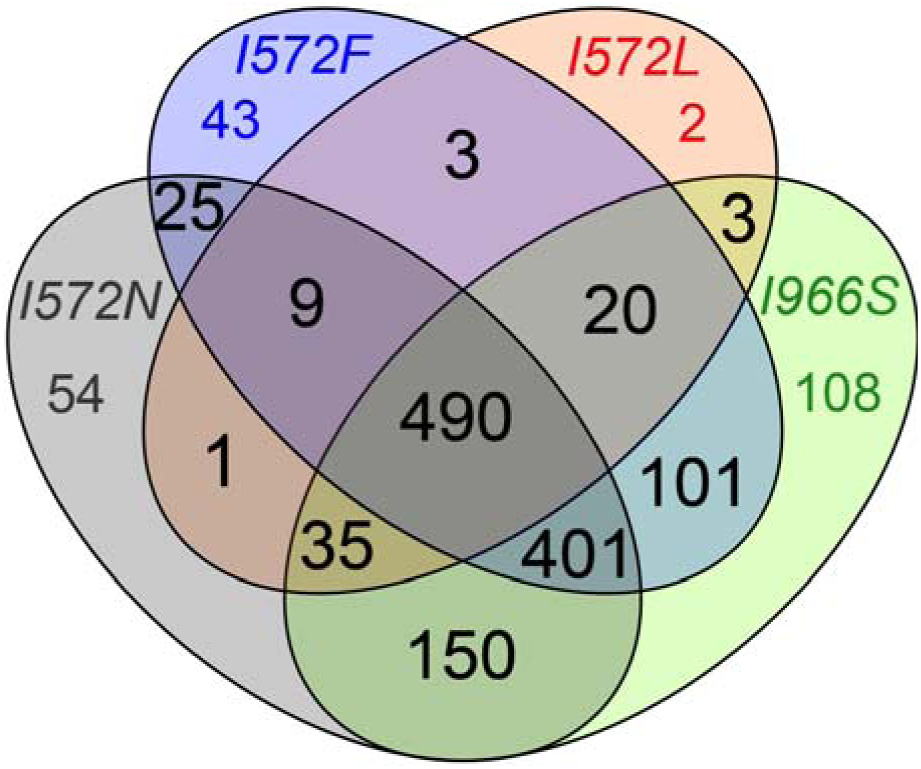
**Convergence of genes with restored expression in single mutants.** Number of shared genes with restored expression.

We also determined how many genes of novel GE were shared between the four single mutants: *I572F*, *I572L*, *I572N* and *I966S*. Only 2% of the genes with novel GE were shared between the four mutants, and 33% of the genes were share by at least two mutants (supplementary fig. S4), indicating lower expression convergence for genes with novel GE than for restored genes.

In conclusion, the phenotypic convergence observed between the mutants *I572F*, *I572L, I572N* and *I966S* suggests: *i*) restoration of the altered physiological state back toward a pre-stressed state, *ii*) that much of that restoration was achieved by single, highly pleiotropic mutations and *iii*) similar effects result from different amino acid mutations at the same site (codon 572 for mutants *I572F*, *I572L* and *I572N*) or at a different site (codon 966 for mutant *I966S*).

### Mutations fixed during adaptation contributed few changes in GE

Following the temporal schematic of this study (fig. 1), our last step was to contrast the GE phenotype of one first-step mutation (*I572L*) to end-products of our adaptation experiment (i.e. clones evolved 2000 generations at 42°C; fig. 6*A*). As end-products, we chose two high-temperature adapted clones: clone 27 and clone 97 (clone numbers correspond to reference Tenaillon et al. 2012), which were isolated from two populations in which the mutation *I572L* swept to fixation before 400 generations (Rodríguez-Verdugo et al. 2013). Although sharing the same *rpoB* mutation, these two high-temperature adapted clones differed in their genetic backgrounds (supplementary tables S4 and S5). Clone 27 had two large deletions of 2,896 and 71,416 bp affecting > 65 genes, a 138 bp deletion disrupting the tRNA-Met gene and seven point mutations in seven genes. In contrast, clone 97 had only one 4 bp small deletion, an IS insertion and eight point mutations in eight different genes (Tenaillon et al. 2012).

**Fig. 6.**
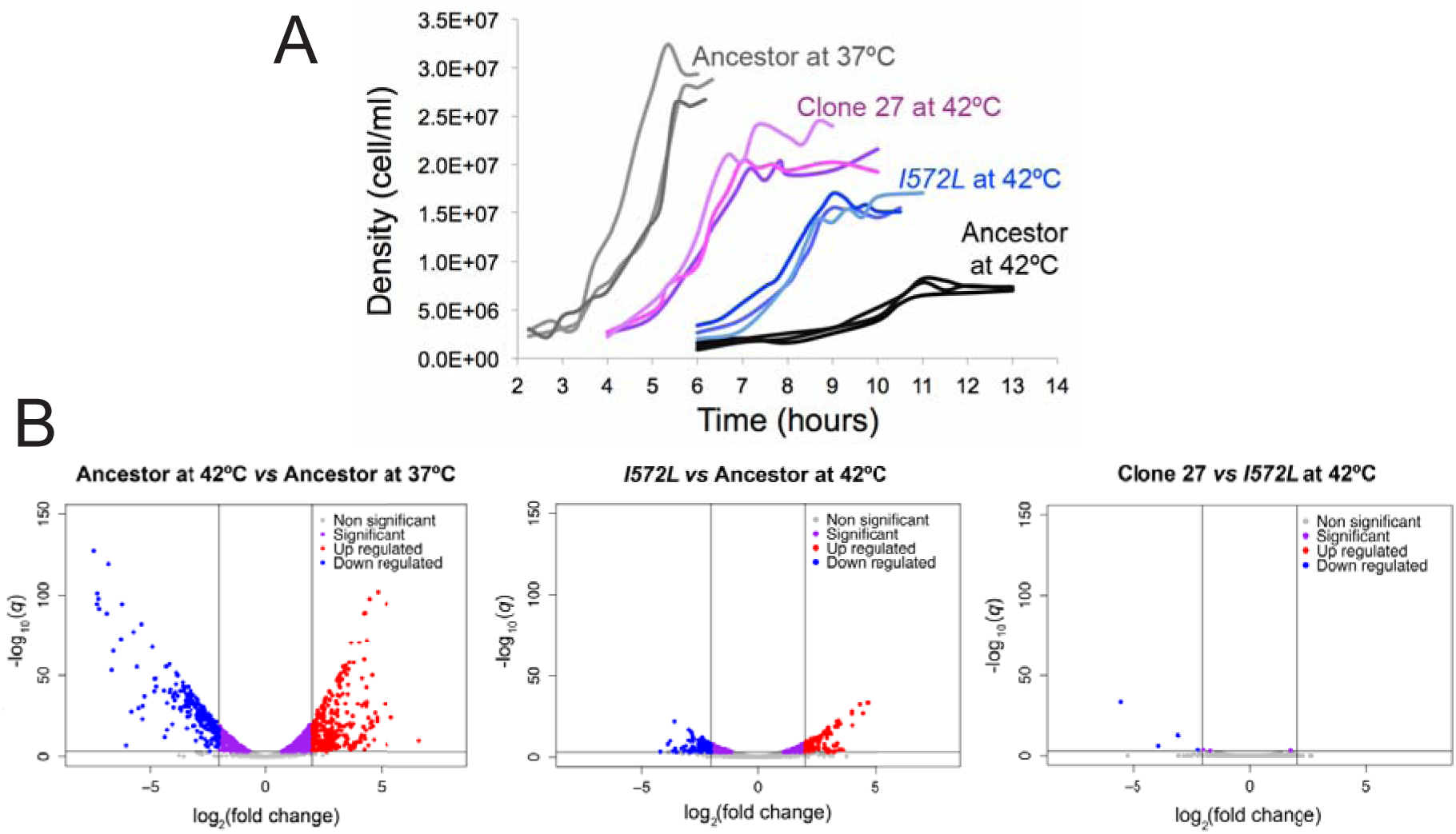
**Phenotypic changes during thermal stress adaptation of clone 27.** (A) Growth improvements during the heat stress adaptive walk. (B) Changes in GE during the acclimation and adaptive response.

We again characterized the clones’ growth curves at 42°C and compared them to both the ancestor grown at 37°C, the ancestor grown at 42°C and the mutant *I572L* grown at 42°C (table 3, fig. 6*A* and supplementary fig. S5). The high-temperature adapted clones 27 and 97 both had significantly shorter lag phases compared to *I572L* mutant. In addition, clone 27 had a significantly higher final yield than *I572L* mutant (table 3). It thus appears that the mutations accumulated after the first-step mutations contribute to better growth at 42°C. We note, however, that none of the adapted clones grow as well as the ancestor at 37°C, which had a significantly shorter lag phase and a higher final yield (table 3 and fig. 6*A*).

**Table 3.** **Growth parameters of the high-temperature adapted clones 27 and 97 at 42°C compare to the mutant *I572L* and the ancestor at 37°C and 42°C (values in table 1).**

| Strain | $\mu_{\max}$ <sup>1</sup> | $\tau$ <sup>2</sup> | Yield <sup>3</sup> |
| --- | --- | --- | --- |
| 27 | 0.769 (0.041) | 4.7 (0.3) | 2.09*10<sup>7</sup> (1.22*10<sup>6</sup>) |
| 97 | 0.794 (0.063) | 4.7 (0.3) | 1.87*10<sup>7</sup> (2.20*10<sup>6</sup>) |
| Comparison | Significance <sup>4</sup> |  |  |
| 27 vs ancestor at 42°C | **0.003** | 0.076 | **0.007** |
| 97 vs ancestor at 42°C | **0.013** | 0.076 | **0.034** |
| 27 vs *I572L* at 42°C | 0.716 | **0.011** | **0.039** |
| 97 vs *I572L* at 42°C | 0.651 | **0.011** | 0.302 |
| 27 vs ancestor at 37°C | 0.801 | **0.049** | **0.011** |
| 97 vs ancestor at 37°C | 0.613 | **0.049** | **0.040** |

<sup>1</sup> Maximum growth rate with standard error in parenthesis.
<sup>2</sup> Duration of the lag phase with standard error in parenthesis.
<sup>3</sup> Final yield (cells/ml) with standard error in parenthesis.
<sup>4</sup> Significance value from a two-sample t-test. The null hypothesis is that the means are equal.

Knowing that the evolved clones grew more quickly and to higher yields than the single mutant *I572L* at 42°C, we sought to identify GE changes that could explain their improved growth. We obtained RNA-seq data from each high-temperature adapted clone (two replicates per clone) grown at 42°C. We then contrasted the GE from each clone against the GE of the first-step mutation *I572L* at 42°C. To our surprise, we observed very few genes that were differentially expressed (*q* <0.001): 7 for clone 27 in addition to 56 genes with no measurable GE (fig. 6*B*, supplementary table S4) and 16 for clone 97 (supplementary fig. S5). Furthermore, the high-temperature adapted clones maintained the general pattern of restoration back to the ancestral physiological state previously observed for the mutant *I572L* (slope of -0.654 for clone 27 and slope of -0.645 for clone 97; supplementary fig. S6).

These observations suggest that the mutations accumulated in later steps of thermal stress adaptation did not substantially change the expression profile caused by the first-step mutation; that is, most detectable GE shifts were caused by the *I572L* mutation and were associated to the restoration of the growth profile. To emphasize this point, we plotted the log_2_fold expression change during the first-step adaptive response (mutant *I572L* grown at 42°C *vs* ancestor grown at 42°C; supplementary fig. S7) against the log_2_fold expression change during the complete adaptive response (high-temperature adapted clone grown at 42°C *vs* ancestor grown at 42°C; supplementary fig. S7). We observed a highly significant positive correlation (slope of 0.977 for clone 27 and slope of 1.004 for clone 97; supplementary fig. S7*A-B*), confirming high similarity in the overall GE pattern between the mutant *I572L* and the high-temperature adapted clones. Finally, when we contrasted the GE from the two high-temperature adapted clones at 42°C (clone 97 *vs* clone 27), there were only 15 differentially expressed genes (*q* <0.001; supplementary fig. S7*C* and fig. S3), beyond the 52 genes that were part of the two large deletions in clone 27 and did not have any measureable GE. Therefore the GE profiles for the two high-temperature adapted clones were nearly identical within the limitations of our experiment, despite their differences in genetic background. These observations confirm that the first step mutation *I572L* contributed to most of the changes in GE during high-temperature adaptation.

## Discussion

Two aspects of adaptation that have been largely unexplored are the temporal change of phenotypes during the adaptive process and the genetic changes underlying these changes. Here we have focused on the phenotypic effects of first-step mutations during adaptation of *E.coli* to high temperature (42°C). A significant finding of our study is that single mutations in RNAP led to altered expression of thousands of genes – most of which were differentially expressed during acclimation to 42°C – and conferred large fitness advantages. A major phenotypic effect of these first-step mutations was to restore global gene expression back towards the pre-stressed state. The GE profile of the ancestor at 42°C revealed that the cells were physiologically stressed, but the first-step mutations restored GE toward an efficient growth profile. Subsequent mutations also increased fitness but did not substantially change GE.

### Heat-stress acclimation involves a balance between energy conservation and stress resistance

The acute response to thermal stress, known as the heat-shock response, occurs in diverse organisms (Richter et al. 2010). In *E.coli,* the heat-shock response is transient (on the order of magnitude of minutes) and is characterized by the induction of stress related proteins mediated by the alternative σ^32^ factor (Nonaka et al. 2006). The σ^32^ regulon encodes: *i)* molecular chaperons (i.e. ClpB, DnaK, DnaJ, IbpB, GrpE, GroEL, GroES) that promote protein folding; *ii)* cytosolic proteases (i.e. ClpP, ClpX) that clear misfolded and aggregated proteins; *iii)* metabolic enzymes; iv) DNA/RNA repair enzymes; *v)* regulatory proteins; *vi)* proteins involved in maintaining cellular integrity; and *vii)* proteins involved in transport and detoxification (Nonaka et al. 2006; Richter et al. 2010). Although the heat-shock response has been studied widely, the expression of heat-shock genes after hours or days of thermal stress (i.e. thermal acclimation) is an open question.

We have found that most heat-shock induced genes (Nonaka et al. 2006; Richter et al. 2010) were not differentially expressed during acclimation and were, in fact, down regulated. For example, most of the heat-shock genes encoding chaperones, such as *clpB*, *dnaJ*, *dnaK*, *groEL* and *groES*, were down regulated during the acclimation response (supplementary table S2). However, one exception is *spy*, which encodes a periplasmic chaperone and was up regulated during acclimation (log_2_fold change = 5.0; *q* < 0.001; supplementary dataset S1). Previous studies in *Saccharomyces cerevisiae* and *E.coli* have reported a decrease in the expression of molecular chaperones after ∼15 minutes of heat-stress induction (Eisen et al. 1998; Zhao et al. 2005; Jozefczuk et al. 2010; Richter et al. 2010). Therefore, it is possible that the heat-shock genes were initially expressed during our experiment, immediately after transfer to 42°C but were subsequently down-regulated.

Nevertheless, two previous studies have explored the physiological acclimation of *E.coli* to high temperature and reported up-regulation of heat-shock genes (Gunasekera et al. 2008; Sandberg et al. 2014). The discrepancy among studies may be explained by differences in genetic backgrounds (*E.coli* K-12 strain *vs E.coli* B strain), by differences in temperature (42°C vs 43°C in Gunasekera et al. 2008), by differences in culture conditions (ours favor microaerophiles) and by differences in sampling. For example, in our study bacterial clones were allowed to acclimate at 42°C for one day before being sampled the next day during mid-exponential phase at 42°C. Thus, the bacteria spent one complete cycle of growth (lag, exponential and stationary phase) at 42°C, in addition to the ∼ half cycle of growth (lag and half exponential phase) at 42°C (in total ∼ 1.5 days). For other studies, the time that the bacteria spent at high temperature before the RNA extraction is unclear (Sandberg et al. 2014). Similarly, we have carefully documented the growth curve of the ancestor at 42°C to help ensure that our RNAseq samples originated from the exponential growth phase (supplementary fig. S8). Other studies report to have sampled during mid-exponential phase but have not reported growth curves. Under normal conditions growth curves may not be necessary, but the exponential phase can be very short under extreme stress. It is thus possible that previous studies have reported gene expression on slightly different phases of the growth cycle.

Microorganisms often resist stressful conditions by modulating GE to limit growth (López-Maury et al. 2008). As a consequence, genes with growth-related functions, which are energy demanding, are down-regulated, allowing a redistribution of resources and energy to the expression of genes related to stress resistance (López-Maury et al. 2008; Jozefczuk et al. 2010; Jin et al. 2012). Surprisingly, we have observed a down-regulation of genes encoding different subunits of RNAP during the acclimation response. Assuming that lower expression also reflects protein abundance, this observation implies that the ancestor contains fewer RNAP molecules when grown at 42°C than when grown at 37°C.

The reduction of RNAP molecules can have important physiological consequences, given that RNAP is limiting for genome-wide transcription (Ishihama 2000). A reduction in the total number of RNAP would limit the transcription rate and favors our hypothesis that mutated RNAP have higher transcriptional efficiency at 42°C (Rodríguez-Verdugo et al. 2014). Accordingly, we have examined RNAP efficiency and found a slight trend toward higher rates of transcriptional efficiency for two of the three mutations in codon 572 of *rpoB*. However, none of these differences were supported statistically (supplementary fig. S1). A limitation in RNAP molecules could also indirectly affect bacterial growth (Jin et al. 2012). For example, when growth conditions are unfavorable, RNAP molecules are released from the rRNA operons, thereby reducing rRNA synthesis (i.e. reducing growth) so that more RNAP molecules become free and available for genome-wide transcription (Jin et al. 2012). Therefore, the reduction in growth at 42°C may be explained in part by the down-regulation of genes involved in translation and ribosome biogenesis but also by a potential limitation of RNAP.

Other down-regulated genes involved in energy demanding processes include genes associated with the biosynthesis of amino acids, nucleotides, ribonucleotides and constituents of the flagella (supplementary table S1). A similar pattern of expression has been observed during acute exposure to thermal stress (Jozefczuk et al. 2010). In addition, metabolomic studies have reported a sharp decline in the levels of nucleotides in *E.coli* cultures exposed to heat stress (Jozefczuk et al. 2010; Ye et al. 2012). Therefore, our study suggests that the pattern of energy conservation mediated through down-regulation of energy demanding processes not only occurs during the acute response to stress but also occurs during the acclimation response.

We hypothesize that some of the resources and energy are redistributed to express genes controlled by the sigma S factor or σ^S^, the master regulator of the stress response (Battesti et al. 2011). Using previous observations (Weber et al. 2005; Keseler et al. 2013), we generated a list of 66 genes expressed under several stress conditions (including high temperature) and under regulation of σ^S^. Of these, 62 genes (94%) were significantly up-regulated during acclimation to 42°C (supplementary table S2). σ^S^ induced genes were related to: *i*) the synthesis molecules responsible to deal with the detrimental effects of stress, *ii*) transport systems, and *iii*) the production of metabolic enzymes, mostly involved in the central energy metabolism (Weber et al. 2005).

In conclusion, the ancestor acclimates to thermal stress by launching alternative GE programs (López-Maury et al. 2008). One involves the down-regulation of growth-related pathways, potentially resulting in energy conservation. The other involves the up-regulation of stress genes that are involved in repair and metabolic adjustments to high temperature.

### Restoration of the ancestral physiological state is advantageous

This study has shown that first-step mutations dramatically altered GE from the acclimated state and acted primarily to restore GE toward the pre-stressed state (fig. 3). Previous studies have also reported that small deletions in RNAP change GE of hundreds of genes (Conrad et al. 2010). Taken together, these observations suggest that *rpoB* mutations have important downstream effects that affect complex networks of interacting genes and their products (i.e. global regulators of GE or “hubs”; (Barabasi and Oltvai 2004)). Our study also adds to previous observations that global regulators are a primary target of natural selection in microbial evolution experiments (Philippe et al. 2007; Hindre et al. 2012).

At least two pieces of evidence suggest that the restoration of GE was advantageous in our experiment, leading to the rapid fixation of *rpoB* mutations in evolved populations (Rodríguez-Verdugo et al. 2013). First, restoration in GE may be advantageous because it “reactivates” some of the growth-related functions that were down-regulated during the acclimation response. The GO categories related to growth, such as translation and ribosomal biogenesis, were significantly enriched in the four mutants relative to the acclimated state. The up-regulation of growth-related genes may explain why the mutants have higher maximum growth rates and higher final yields than the ancestor (fig. 3), as well as higher relative fitness values (Rodríguez-Verdugo et al. 2013; Rodríguez-Verdugo et al. 2014)).

Second, the general trend of restoration occurred in parallel in all four *rpo*B mutants (*I572F*, *I572L, I572N* and *I966S*); such phenotypic convergence is commonly interpreted as sign of adaptive evolution (Christin et al. 2010). Not only did we observe convergence in global GE (fig. 4), but we also observed substantial overlap of genes with restored expression (fig. 5). Surprisingly, these observations extend not only to three different mutations in the *rpoB* codon (572) but also to a mutation (*I966S*) that is not in the RNAP active site (fig. 4*D* and fig. 5). It is unclear whether the effects of these RNAP mutants are direct or indirect. For a direct effect, the mutated RNAPs would have had new intrinsic binding affinities for promoters throughout the genome. Alternatively because RNAP is a sensor of the physiological state of the cell (Ishihama 2000), it may induce different GE programs. For example, under amino acid starvation RNAP modifies recruitment of sigma factors through its interaction with the alarmone ppGpp, and it consequently induces the stringent response program (Chatterji and Ojha 2001). The fact that mutations with similar effects are not limited to the active site of the RNAP favors the possibility of indirect effects, but further experiments are needed to test this hypothesis.

It is worth noting that restoration to the non-stress phenotype is not complete, because first-step mutations differ markedly from the 37°C ancestor in GE dynamics. For example, each of the four *rpoB* mutations differ from the ancestor in the expression of a minimum of 392 genes (*I572L*) and as many as 688 genes (*I572N*). The important point is that first-step mutations restore hundreds of genes toward non-stress expression dynamics but substantial differences remain relative to the wild type phenotype. These differences may explain, in part, differences in growth characteristics between first-step mutations and the ancestor at 37°C (supplementary fig. S9).

In contrast to restorative changes in GE, we find that few novel changes in GE occurred in parallel among the four *rpo*B mutants (supplementary fig. S4). These observations add to a growing literature that suggests restorative changes in GE are a general trend in laboratory adaptation experiments, but novel (and/or reinforced) changes are less common (Fong et al. 2005; Carroll and Marx 2013)(Sandberg et al. 2014); although see (Szamecz et al. 2014).

### Pleiotropy and compensation

The four *rpoB* mutations (*I572F*, *I572L*, *I572N* and *I966S*) were advantageous in the conditions of our experiment, but they also had pleiotropic effects, such as fitness trade-offs at low temperatures (Rodríguez-Verdugo et al. 2014). Previous studies have shown that mutations in global regulators often have maladaptive side effects (Hindre et al. 2012). We hypothesized, therefore, that later adaptive mutations compensate for maladaptive side effects of highly pleiotropic first-step mutations (Hindre et al. 2012).

To contrast first-step mutations to the end-products of our adaptation experiment (fig. 1), we compared the *I572L rpoB* mutation to two clones (97 and 27) that included the *I572L* mutation. Each of the two clones differed from the single-mutant background by ∼10 mutations; these mutations included both point mutations and more complex genetic changes, such as a large deletion. Based on these contrasts, we have found that the first-step mutation *I572L* contributed most of the GE variation during thermal stress adaptation (987 differentially expressed genes), while later mutations contributed fewer changes in GE (63 and 16 differentially expressed genes in clones 27 and 97, respectively; fig. 6 and supplementary fig. S5). Interestingly, these few changes in GE may contribute to significant changes in growth parameters (table 3). Therefore, the number of differentially expressed genes was not proportional to fitness gains.

This “disconnect” between the number of genes differentially expressed and the magnitude of the fitness advantage might be caused by compensatory changes to the pleiotropic side effects of mutation *I572L*. Under this model, we presume that some of the hundred of differentially expressed genes in *I572L* have beneficial effects, while others have deleterious effects, netting an overall beneficial change in GE. If true, it is likely that subsequent mutations change the expression of only few genes, but most of these changes are beneficial. For example, the large deletion in clone 27 contained genes involved in iron acquisition (*fep* and *ent* operons; Crosa and Walsh 2002), and copper and silver efflux system (*cus* operons; Long et al. 2010). Costly, non-functional pathways are often “shut-down” in order to save energy that would be otherwise used to produce unnecessary proteins and metabolisms (Cooper et al. 2001; Lewis et al. 2010). Therefore the large deletion of 71 kb in length may be an energetic benefit, explaining its occurrence in 35 high-temperature adapted clones (Tenaillon et al. 2012).

Clone 97, which lacked the large deletions of clone 27, displayed fewer changes in gene expression than clone 27. In this clone, one of the significantly down-regulated genes (*q* < 0.001) was the *rmf* gene, which encodes the ribosome modulation factor (RMF). RMF has been associated with decreased translation activity and is expressed during slow growth conditions, such as stationary phase (Polikanov et al. 2012). Therefore, the down-regulation of *rmf*, occurring in parallel in clone 27 and 97, might increase protein synthesis and thus growth.

Finally, we observed that the genes involved in flagellum-dependent cell motility (e.g. *flg* genes) were highly down-regulated during the acclimation response (log_2_fold < -3 and *q* < 0.001) but were again up-regulated in the single mutants *I572F* and *I572L* (*q* < 0.001; supplementary dataset S1). In the conditions of our evolution experiment — a well-mixed environment lacking physical structure — motility seems unnecessary. Given that the biosynthesis of flagella is costly (Soutourina and Bertin 2003), reducing the expression *flg* genes might be beneficial (Cooper et al. 2003; Fong et al. 2005). We therefore posit that the restoration in GE of *flg* genes might be costly and have deleterious effects. Interestingly, some of the *flg* genes were down-regulated (*q* < 0.05) in clone 97 when compared to the first-step mutant *I572L* (supplementary table S6). Therefore, later adaptive mutations might contribute to the fine-tuning of GE by compensating the side effects of restoration. We note, however, that we have not yet identified the mutation that causes the down-regulation of *flg* genes in clone 97. That being said, the up-regulation of flagellar genes after restorative shifts in GE and the fixation of subsequent mutations that down-regulate them has been observed previously (Sandberg et al. 2014).

### Concluding remarks

Mutations in global regulators of GE are observed recurrently in laboratory evolution experiments (Applebee et al. 2008; Goodarzi et al. 2009; Kishimoto et al. 2010). It is not always clear if these mutations represent the first step of an adaptive walk or later step, but at least in the case of *rpoB* and *rpoC* mutations, it seems they are often first-step mutations (Herring et al. 2006). Therefore, the pattern that we have observed in our study may not be specific to our system but instead a more general phenomenon. Based on our results, we propose a general, two-step adaptive process. First, a mutation affecting global transcriptional regulator appears in the population and changes global GE. This expression change is mostly restorative, so that the stressed state moves toward a pre-stressed (ancestral) state. These changes in GE confer a high advantage, promoting the rapid fixation of the mutation in the population. Once the cell recovers its “normal” homeostatic state, other mutations accumulate and contribute to novel functions (fine-tuning of adaptive traits) or/and compensate for the side effects of the first-step pleiotropic mutation. Future directions would be to confirm this pattern by performing time course studies of GE (including heat shock and acclimation response, as well as all the intermediate steps of adaptation) coupled with genomic data.

## Materials and Methods

### Growth conditions

Unless otherwise noted, the culture conditions used for the physiological assays (growth curves, transcription efficiency and RNA-seq assays), were the same used during the high-temperature evolution experiment (Tenaillon et al. 2012). Briefly, strains were revived in LB and incubated at 37°C with constant shaking (120 rpm). Overnight cultures were diluted 10^4^-fold into 10 ml Davis minimal medium supplemented with 25 μg/ml glucose (DM25) and incubated 1 d at 37°C to allow the strains to acclimate to the culture conditions. The following day, we transferred 100 μl of the overnight culture in 9.9 ml of fresh DM25 (100-fold dilution) and we incubated them at 42°C for one day to allow the strains to acclimate to high temperature.

### Growth curves

Acclimated strains were grown at the assay temperature (either 37°C or 42°C) and the densities were measured approximately every hour during the lag phase and every ∼30 minutes during the exponential phase. Population densities were measured using an electronic particle counter (Coulter Counter model Multisizer 3 equipped with a 30-μm-diameter aperture tube). To measure density, 50 μl of culture was diluted in 9.9 mL Isoton II diluent (Beckman Coulter), and 50 μl of the resulting dilution was counted electronically. To estimate the maximum growth rate, μ_max_, we fitted a linear regression to the natural logarithm of the cell density over time during the exponential phase (i.e. linear part) using the *lm* function in R version 3.0.2 (Team 2013, supplementary fig. S8). Three estimates of μ_max_ were obtained for each genotype. The duration of the lag phase, τ, was defined as the time elapsed before the beginning of the exponential phase. The final yield was estimated from the cell counts at the end of the exponential phase.

### RNA extraction and preparation for sequencing

To investigate the GE profile prior to thermal stress, we grew the ancestor (REL1206), previously acclimated to the growth conditions, at 37°C until it reached mid-exponential phase (three replicates). To investigate the GE profile during acclimation, we grew the ancestor at 42°, previously acclimated to 42°C (two replicates), until mid-exponential phase. The remaining strains (single first-step mutants and high-temperature adapted clones) were grown at 42°C until mid-exponential phase, with two replicates for each.

Briefly, bacterial cultures were grown in DM25 medium until they reached mid-exponential phase, which we determined by electronic counts. 80 ml of culture was filtered through a 0.2 μm cellulose membrane (Life Science, Germany). Cells, concentrated in the filter, were stabilized with Qiagen RNA-protect Bacteria Reagent and pellet for storage at -80°C prior to RNA extraction. Total RNA was extracted using RNeasy Mini Kit (Qiagen). Total RNA was DNase treated using Turbo DNA-free kit (Ambion) and rRNA was depleted using the Ribo-Zero rRNA Removal kit for Gram-Negative Bacteria (Epicentre Biotechnologies, Medion, WI, USA). cDNA library was constructed using TruSeq RNA v2 kit (Illumina, San Diego, CA, USA). Libraries were multiplexed 8-fold and sequenced on an Illumina Hiseq 2000 platform. 100-bp single-end reads were generated.

### mRNAseq data analyses

Reads were mapped to the *E.coli* B REL606 genome reference (CP000819.1) using *bwa* 0.6.2 (Li and Durbin 2009), using default parameters (http://bio-bwa.sourceforge.net/bwa.shtml). Only unique, perfectly matching reads to the 4204 annotated coding regions were retained for further analyses (supplementary table S7). Differential expression analysis was performed using the DESeq R package (Anders and Huber 2010). We used the *P*-values adjusted by the Benjamini and Hochberg approach (*q* values), which controls for false discovery rate. Genes with *q* less than 0.001 were considered significantly differentially expressed. For the GO term enrichment analyses, we used the Enrichment analysis tool from http://geneontology.org/page/go-enrichment-analysis.

Differentially expressed genes (DEG) were classified in one of the four categories (restored, reinforced, unrestored, and novel) based on the contrasts for which they were significant and the direction and value of their fold change (supplementary table S3). We also examined overall trends in GE shifts by plotting two ratios that share an underlying parameter (fig. 4 and supplementary fig. S6), which creates an inherent negative bias for correlation values. We therefore tested the strength of correlation by randomization; we choice the three independent parameters at random from the dataset, constructed ratios for the observed number of genes, estimated a correlation coefficient, and repeated the process 1000 times to achieve a distribution of correlation values that reflect the inherent bias.

### Construction of the fluorescently labeled strains for the transcription efficiency assay

We generated a ∼4 kb-long linear DNA fragment carrying the *CAT*, *tetR* and *YFP* genes from an *E.coli* strain carrying the *CAT:tetR:YFP* genomic cassette (Fehér et al. 2012) kindly provided by Csaba Pál (Biological Research Center, Szeged, Hungary). We amplified the *CAT;tetR;YFP* cassette by PCR using the primers ARV19 and ARV20 (supplementary table S8) and *Pfu* DNA polymerase (Promega). The purified PCR product was integrated into the ancestral strain REL1206 carrying the pKD46 recombineering plasmid as previously described by Datsenko and Wanner (Datsenko and Wanner 2000). Briefly, the ancestral strain carrying the pKD46 plasmid was grown overnight at 30°C in 5 ml of LB with 100 μg/ml of ampicillin. The overnight culture was 100 fold-diluted in 100 ml of LB with ampicillin and 1 mM L-arabinose (Sigma) and grown at 30°C to an OD_600_ of 0.6. Electrocompetent cells were made by washing the culture 5 times with ice-cold water. ∼200 ng of linear DNA was electroporated into 25 μl of cells. After electroporation, 1 ml of LB was added, and the cells were incubated at 30°C for 2 h with shaking, then plated 100 μl on LB agar plates with chloramphenicol (20 μg/ml). We selected a single colony and purified it in LB agar plate containing chloramphenicol. Correct integration was verified by PCR using primers ARV34 and ARV35 (supplementary table S8).

### Transcription efficiency assay

To measure the transcription efficiency of the ancestor and the mutants we performed a quantitative reverse transcriptase RT-PCR assay (Reynolds 2000; Brandis et al. 2012). In brief, we grew the fluorescently labeled strains in the same culture conditions previously described except that we supplemented the DM medium with 100 μg/ml glucose (DM100). We confirmed the advantage of these mutations to high temperature despite the higher amount of glucose based on growth curves of the ancestor and the mutants at 42°C in DM100. Acclimated cultures were grown at 42.2°C until they reached mid-exponential phase. We took 1 ml of uninduced cells (sample T_0_) and stabilized them in RNAprotect Bacteria Reagent (Qiagen). Immediately after, we induced cells by adding 10μl of anhydotetracycline (66 μg/ml) to the medium. Samples of 1 ml of culture were taken at 1, 2, 3 and 4 min after induction (samples T_1_, T_2_, T_3_ and T_4_) and stabilized with RNA protect Bacteria Reagent and pelleted for storage at - 80°C prior to RNA extraction. Total RNA was prepared using RNeasy Mini Kit (Qiagen). RNA was DNase treated using Turbo DNA-free kit (Ambion). We used 300 ng of DNA-free RNA to produce cDNA with the High Capacity cDNA Reverse Transcription Kit (Applied Biosystems). From each reverse transcribed product we quantified the abundance of cDNA of the gene reference *gst* (Gst, glutathione transferase; Pfaffl 2001) and the target gene YFP (Yellow fluorescent protein) using Fast SYBR Green Master Mix (Applied Biosystems) quantified on a Stratagene MX3005P QPCR System (Agilent Technologies). For each PCR reaction we used 0.625 μM forward primer (ARV48 for *gst* or ARV50 for YFP, supplementary table S8), 0.625 μM reverse primer (ARV49 for *gst* or ARV51 for YFP, supplementary table S8), 3 μl CDNA template, 4.5 μl RNase-free water and 10 μl of Fast SYBR Green Master Mix, to have a final reaction volume of 20 μl. To control for the intra-assay variation (repeatability), we prepared three replicates of each reaction. The PCR thermal cycling conditions were 95°C for 20 sec followed by 40 cycles of 95°C for 3 sec and 60°C for 30 sec.

The efficiency of the amplifications for each pair of primers was determined from a standard curve using the formula E = 10^[-1/s]^, where *s* is the slope of the standard curve (Pfaffl 2001). To calculate the *Relative Expression Ratio* (i.e. the relative change in GE of the target gene YFP normalized to the reference gene *gst* and relative to the uninduced control sample T_0_), we used the mathematical model for relative quantification in real-time RT-PCR developed by Pfaffl, 2001 (Pfaffl 2001; Brandis et al. 2012). Transcription efficiency was calculated as the slope of the fitted linear regression between the *Relative Expression Ratio* against time, based on three replicates.

## Acknowledgments

We thank P. McDonald and R. Gaut for technical assistance. This work was supported by National Science Foundation Grant DEB-0748903 to BG. OT was supported by European Research Council under the European Union’s Seventh Framework Programme (FP7/2007-2013)/ERC Grant 310944. AR-V was supported by University of California Institute for Mexico and the United States-Consejo Nacional de Ciencia y Technología (Mexico) Fellowship and by the Chateaubriand Fellowship in Science, Technology Engineering, and Mathematics (STEM).

